# Raabin-WBC: a large free access dataset of white blood cells from normal peripheral blood

**DOI:** 10.1101/2021.05.02.442287

**Authors:** Zahra Mousavi Kouzehkanan, Sepehr Saghari, Eslam Tavakoli, Peyman Rostami, Mohammadjavad Abaszadeh, Farzaneh Mirzadeh, Esmaeil Shahabi Satlsar, Maryam Gheidishahran, Fatemeh Gorgi, Saeed Mohammadi, Reshad Hosseini

**Affiliations:** School of ECE, College of Engineering, University of Tehran, Tehran, Iran; Nimaad Health Equipment Development Company, Tehran, Iran; Graduated Bachelor of laboratory of sciences, Paramedical Faculty of Guilan University of Medical of Sciences, Langroud, Guilan, Iran; Faculty of Electrical Engineering, K. N. Toosi University of Technology, Tehran, Iran; Master Graduated, School of Mechanical Engineering, Sharif University of Technology, Tehran, Iran; Graduated from School of ECE, College of Engineering University of Tehran, Tehran, Iran; M.A Medical Biotechnology, Graduated from Tarbiat modares University, school of medicine, Tehran, Iran; Ph.D of Hematology, Takhte tavous patobiology lab, flow cytometry department, Tehran, Iran; M.A Hematology, Department of Hematology and Blood Transfusion, School of Allied Medical Sciences, Iran University of Medical Sciences, Tehran, Iran; Bachelor of labrotoary of Sciences, Mashhad University of Medical Sciences, Faculty of Paramedical, Mashhad, Iran; Hematology-Oncology and Stem Cell Transplantation Research Center, Tehran University of Medical Sciences, Tehran, Iran

**Author notes:** Equally Contribution (Eslam Tavakoli, Peyman Rostami, Mohammadjavad Abaszadeh).

## Abstract

Accurate and early detection of peripheral white blood cell anomalies plays a crucial role in the evaluation of an individual’s well-being. The emergence of new technologies such as artificial intelligence can be very effective in achieving this. In this regard, most of the state-of-the-art methods use deep neural networks. Data can significantly influence the performance and generalization power of machine learning approaches, especially deep neural networks. To that end, we collected a large free available dataset of white blood cells from normal peripheral blood samples called Raabin-WBC. Our dataset contains about 40000 white blood cells and artifacts (color spots). To reassure correct data, a significant number of cells were labeled by two experts, and the ground truth of nucleus and cytoplasm were extracted by experts for some cells (about 1145), as well. To provide the necessary diversity, various smears have been imaged. Hence, two different cameras and two different microscopes were used. The Raabin-WBC dataset can be used for different machine learning tasks such as classification, detection, segmentation, and localization. We also did some primary deep learning experiments on Raabin-WBC, and we showed how the generalization power of machine learning methods, especially deep neural networks, was affected by the mentioned diversity.

## Introduction

The issue of precise and early diagnosis is the most important step in the medical treatment process. According to the World Health Organization, about 2 billion people currently do not have access to basic medical and pharmaceutical services [1]. In the meantime, laboratory tests play an essential role in the diagnosis and treatment of the diseases. It is estimated that about 70% of the decisions related to the diagnosis and treatment of the disease, as well as the discharge and admission of a patient, rely on the results of laboratory tests [2]. In this regard, the differential count of white blood cells is one of the common laboratory tests necessary to be considered in various diseases such as blood disorders (such as leukemia, anemia, polycythemia, etc.) and immune system related diseases (such as autoimmune anemias, allergy. etc.) and are of utmost significance [3].

White blood cells called leukocytes fall into two groups of phagocytes and lymphocytes. While phagocytes comprise cells of the innate immune system and function rapidly after infection, lymphocytes mediate the acquired immune response. Phagocytes, themselves, can be divided into granulocytes (neutrophils, basophils, and eosinophils) and monocytes. In Table 1[4] and Figure 1, you can see the characteristics and images of the five categories of white blood cells, respectively. Table 2 [5], also, shows some examples of the diseases that occur with an increase or decrease in the number of white blood cells. For example, in allergic diseases, one of the types of white blood cells (specifically basophils) increases, or in blood malignancies, we can see an increase in the number of precursors of blood cells and changes in their shape and size. Therefore, determining the correct type and number of white blood cells is very important for diagnosing various diseases.

**Table 1.** Characteristics of white blood cells [4].

| WBCs | % in blood | Nucleus | Cytoplasm | Size (μm) |
| --- | --- | --- | --- | --- |
| Neutrophils | 60% | Divided into 2 to 5 segments and stains dark purple (multi-lobed nucleus) | Pale pink to tan with pink-purple granules | 12-16 |
| Eosinophils | 3% | Blue and is divided into 2 segments | Full of pale pink tan with large orange and red granules | 14-16 |
| Basophils | 1% | Have 2 lobes that stains purple and is difficult to see | Pale pink-tan but contains large purple/blue-black granules obscure nucleus | 14-16 |
| Monocytes | 6% | Have singular nucleus (convoluted shape) kidney shaped, bean shaped or horseshoe shaped with deep indentation | Stain a blue-gray color with "ground glass" cytoplasm with tiny granules, Vacuoles are sometimes present. | 14-20 |
| Lymphocytes | 30% | Have large, round or oval, dark staining nucleus. | Little to no cytoplasm with pale blue in color. Occasional purple-reddish granules | 8-15 |

**Table 2.** White blood cells alterations and related different diseases [5].

| White Blood Cell | Increase | Decrease |
| --- | --- | --- |
| Lymphocyte | Acute & Chronic Leukemia, Hypersensitivity reaction, Viral infection | AIDS, Influenza, Sepsis, Aplastic Anemia |
| Monocyte | Autoimmune disease, Fungal and Protozoan infection | Aplastic Anemia, Hairy cell leukemia, Acute infections |
| Neutrophil | Chronic inflammation, Infection | Chediak-Higashi syndrome, Kostmann syndrome, Autoimmune Neutropenia |
| Eosinophil | Allergic reaction, Parasitic infection, Malignancy | Cushing syndrome, Shock or trauma driven stress |
| Basophil | Leukemias | Hyperthyroidism & Acute infections |

**Figure 1.**
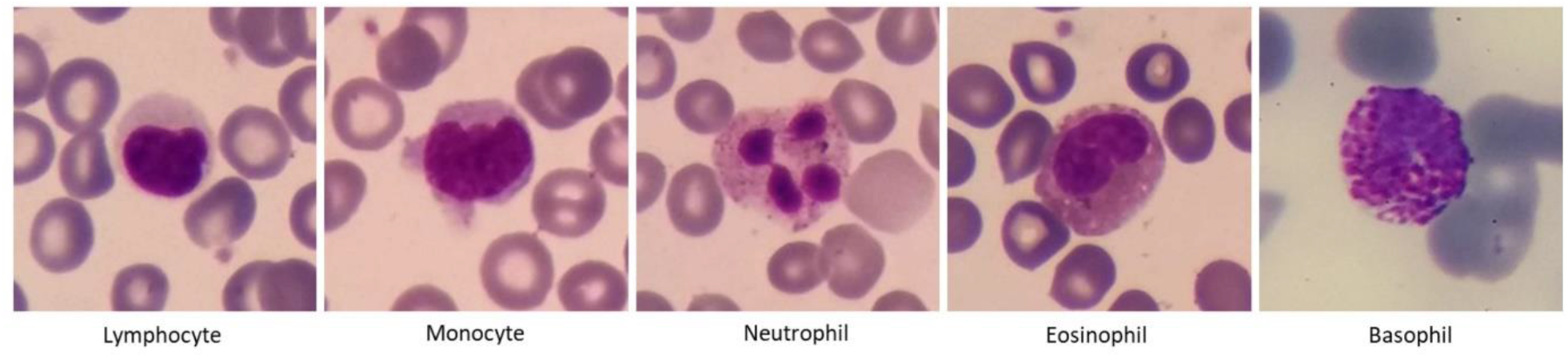
Five types of white blood cells in normal peripheral blood

At present, manual (microscopic evaluation) and automated methods (using automatic hematology devices) are used to evaluate blood cells. Automated methods include devices which evaluate blood cells based on light scattering or electrical impedance such as Sysmex XP-300, Nihon Kohden Blood Cell Counter, and DH36 3-Part Auto Hematology Analyzer.

In electro-optical analyzers, a light-sensing detector measures the optical scattering. The size of the detected pulses corresponds to the size of the blood cells. Furthermore, in electrical impedance or Coulter principle cell counter, the passage of cells through an aperture in which an electric current is applied causes change in electrical resistance and is counted as voltage pulse. Pulses the height of which corresponds to the volume of the cell are counted, and this is considered as the basis of Coulter’s principle working [4].

One of the serious drawbacks of these devices apart from their high cost is the simple act of counting cells without them being evaluated qualitatively from a structural and morphological point of view. As a result, after evaluating the blood sample by the mentioned cell counters, it is necessary to prepare a smear and evaluate it microscopically by the laboratory staff in order to achieve an accurate and correct diagnosis.

On the other hand, issues such as lack of specialists and laboratory equipment, heavy workload, inexperience, and incorrect diagnosis affect the test results. Incorrect diagnosis affects the treatment regime, and consequently, can result in incorrect treatment and increase the associated costs. However, the use of new technologies such as artificial intelligence and image processing allows quantitative and qualitative evaluation to improve the quality of diagnosis [6].

Over the past 20 years, the techniques for automated imaging of the blood-stained slides have been introduced by computer-connected microscopes capable of assessing blood cell morphology. With the development of technology, implementing modern machine learning techniques (such as neural networks) and image processing, companies such as Cellavision, Westmedica, Siemens, etc. have made it possible to differentiate count of normal from abnormal blood cells [7]. In fact, today, deep neural networks are one of the most widely used machine learning methods for classification and segmentation of the medical images. Shahin, A et al. [8] used DNN in order to classify white blood cells. In addition, these networks are used for the classification of the red blood cells to detect a sickle cell anemia [9]. Deep neural networks are also used for segmentation of the pancreas in the CT scan images [10,11] and segmentation of the MRI images [7].

Data have the most important role to play in the development of machine learning models. In order to train deep neural networks and increase their generalizability, we need a lot of diverse precise data and confident labels. The process of labeling medical data should be carried out by professionals and is, therefore, a time consuming and challenging procedure. As a result, medical databases are of high significance in smartening medical diagnoses. Unfortunately, researchers, today, have limited access to a variety of medical data for various reasons. Examples of available medical image databases are [12] and [13]. The database [12] contains 82 3D CT scans in which the Grand Truth of pancreas for all slices were manually extracted by medical students and finalized by a specialist radiologist. Camelyon [13] is another dataset with 1399 whole-slide images (WSIs) of the lymph node smear samples with and without metastases, which was evaluated twice.

The morphological diversity of white blood cells is very high and in some cases, it is very challenging, even for an expert, to distinguish some classes from each other. On the other hand, many artificial intelligence articles have adopted two approaches to evaluate their proposed method regarding segmentation and classification of white blood cells: They have either collected small databases to the best of their ability [14, 15, 16, and 17] or used the small databases available [18, 19, and 20]. Therefore, a database with a large amount of diverse data and reliable labelling is truly necessary to evaluate and compare different methods with each other. Such a reference database will allow more artificial intelligence scientists to enter the field and will help the advancement of intelligence differentiation of white blood cells. The most important characteristics of the Raabin-WBC dataset that distinguishes it from similar datasets are as follows:

- **Large number of data**: We tried to collect as much data as possible for each class in order for them to be appropriate for all machine learning techniques, especially deep learning. (Approximately 40,000 white blood cell images)
- **Precise labels**: We considered more detailed labels than five types of white blood cells. In fact, labels contain the most important subgroup of each type. For example, we considered the meta and band which are subgroups of neutrophils and are valuable in respect of diagnosis. In the next section, more information about the labels will be presented.
- **Double labeling**: For more insurance, most of the cells are labeled by two experts.
- **Free public access**: Since we aim at helping the development of AI in hematology, the Raabin-WBC dataset is freely available for all.
- **Data cleaning**: In the process of data collection, the existence of duplicate cell images is not inevitable. The first problem is that the duplicate cell images are not exactly the same. For example, there is a possibility of the cell being somewhat moved. The second issue is that having more than two versions of one cell image is also possible. Hence, we developed a fast graph-based image processing method that can accurately remove as many duplicate cells as possible. Despite this, it is still probable for some duplicate images to exist, albeit being significantly different.
- **The nucleus and cytoplasm ground truth**: The ground truth of the nucleus and cytoplasm are extracted for 1145 selected cells. In order to extract the nucleus ground truth, we developed a software which uses image processing tricks to make the ground truth extraction process much easier.
- **Diversity of the microscope and camera**: Although most of the data were collected by a fixed type of microscope and camera, we collected some data with another type of camera and microscope, as well. In the section of experiments, you will see how new test data help to evaluate the generalization power of our trained models. In other words, the diversity of the dataset assists us in selecting a model that has correctly learned the manifold of cell images.

The rest of the paper is as follows: In section 2, we will elaborate more on the details regarding the dataset. In section 3, the data collection process will be explained completely. In section 4, we will do some machine learning experiments and discuss the generalization power of the models.

### The characteristics of Raabin-WBC

In this section, more information is provided about the Raabin-WBC dataset. About 73 peripheral blood films were used for collecting this dataset. After imaging stained blood films, we tried to mine the most possible useful information from raw data. For instance, the bounding box of all white blood cells and artifacts were extracted, cropped and labeled, successively. It is worth noting that a significant number of WBCs and artifacts were labeled by two experts. Furthermore, we provided the ground truth of the nucleus and cytoplasm for some of the cropped cells. The full details of the data collection steps are explained in section 3. In table 3, some general and useful information of the Raabin-WBC dataset is provided. Note that these numbers have been computed after cleaning phase.

**Table 3.**
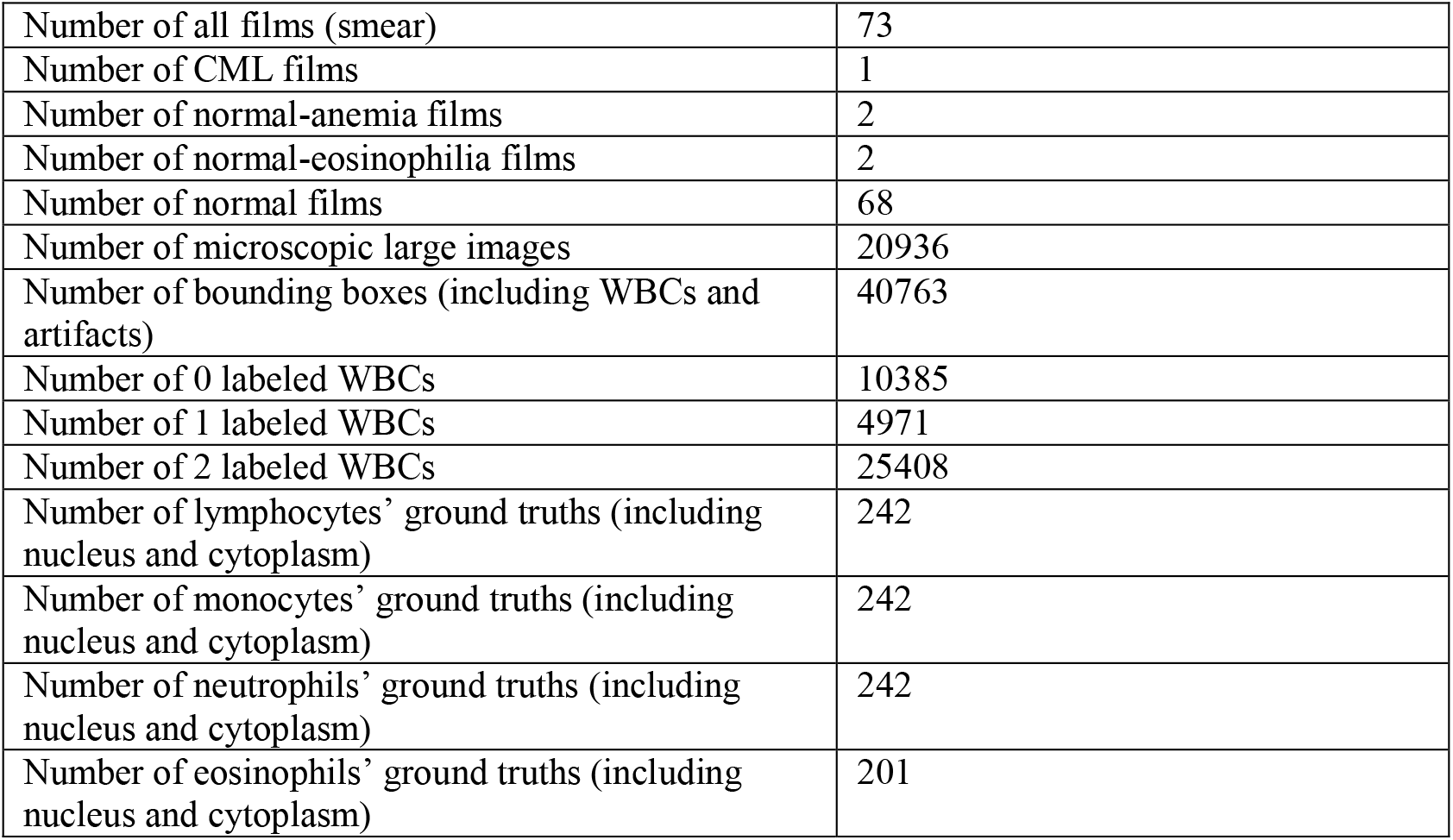
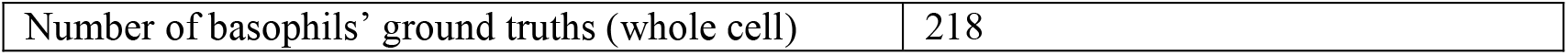
Raabin-WBC information table.

|  |  |
| --- | --- |
| Number of all films (smear) | 73 |
| Number of CML films | 1 |
| Number of normal-anemia films | 2 |
| Number of normal-eosinophilia films | 2 |
| Number of normal films | 68 |
| Number of microscopic large images | 20936 |
| Number of bounding boxes (including WBCs and artifacts) | 40763 |
| Number of 0 labeled WBCs | 10385 |
| Number of 1 labeled WBCs | 4971 |
| Number of 2 labeled WBCs | 25408 |
| Number of lymphocytes' ground truths (including nucleus and cytoplasm) | 242 |
| Number of monocytes' ground truths (including nucleus and cytoplasm) | 242 |
| Number of neutrophils' ground truths (including nucleus and cytoplasm) | 242 |
| Number of eosinophils' ground truths (including nucleus and cytoplasm) | 201 |
| Number of basophils' ground truths (whole cell) | 218 |

#### Labels

In Raabin-WBC dataset, more detailed labels are considered than just five general types of white blood cells. For example, beside the mature neutrophil, we have evaluated two other ancestors of this white blood cell: Metamyelocytes and Band. An increase in the number of band forms and metamyelocytes is one of the features of reactive neutrophilia (an increase in the number of circulating neutrophils to levels greater than 7.5 × 109/L)[5]. In addition, lymphocytes are divided into small (the main agents of the acquired immune system including B & T cells) and or activated lymphocytes (activated small lymphocyte referred to as large lymphocytes or lymphoblasts). The Burst label belongs to smudge cells that are leukocyte remnants formed during blood smear preparation. Beside the leukocytes we considered drying artifacts as new labels for artifacts are commonly seen after staining the samples. In figure 2 the diagram of the labels is presented.

**Figure 2.**
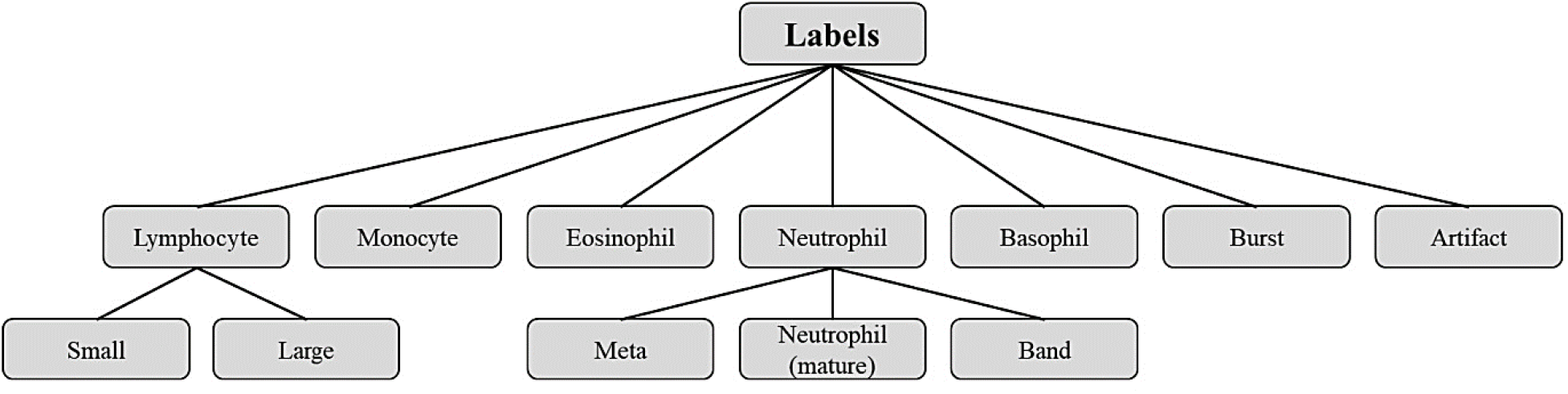
Raabin-wbc dataset labels diagram.

In table 4, the number of labels associated with two experts is shown. The rows and columns of the table belong to the first and second experts, respectively, noting that 9015 cells have not been labeled yet. We asked our experts to label the cells as unrecognizable if they had any doubts. Indeed, we have 1099 cells labeled as not recognized by the two experts. In table 4, you can see the amount of disagreement for each pair of different labels (Non-diagonal elements of the matrix). For example, large and small lymphocytes are confused a lot. Also, seem bands have often been mistaken with mature neutrophils. Other examples of confusing pairs are Artifact and burst, large lymphocyte and monocyte, and small lymphocyte and burst. The high numbers in the rows and columns labeled as not recognized indicate that it is very challenging to identify the type of white blood cell.

**Table 4.**
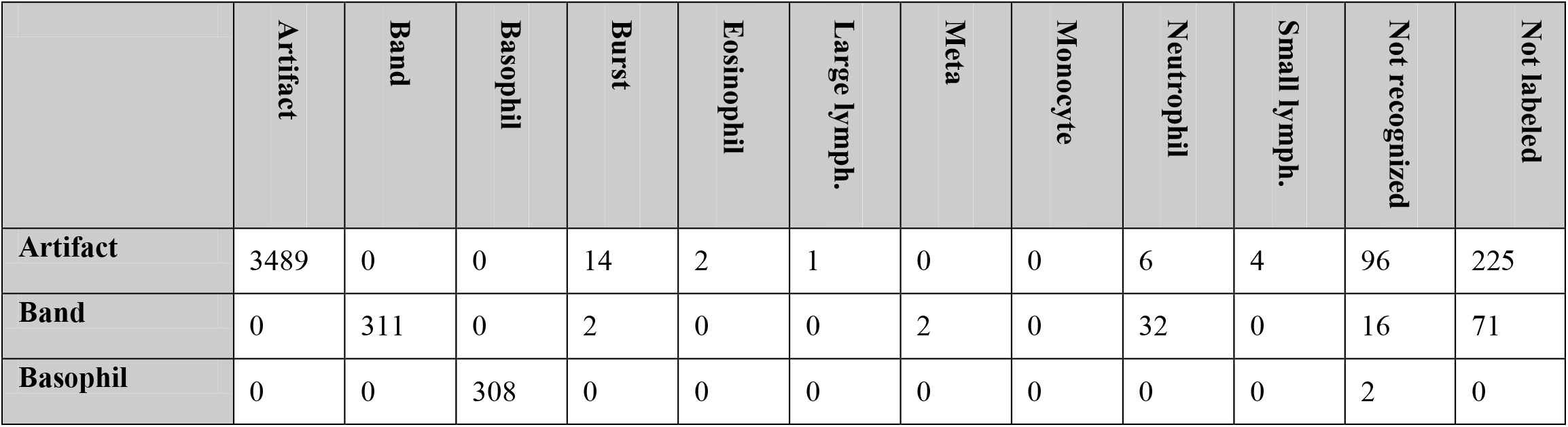
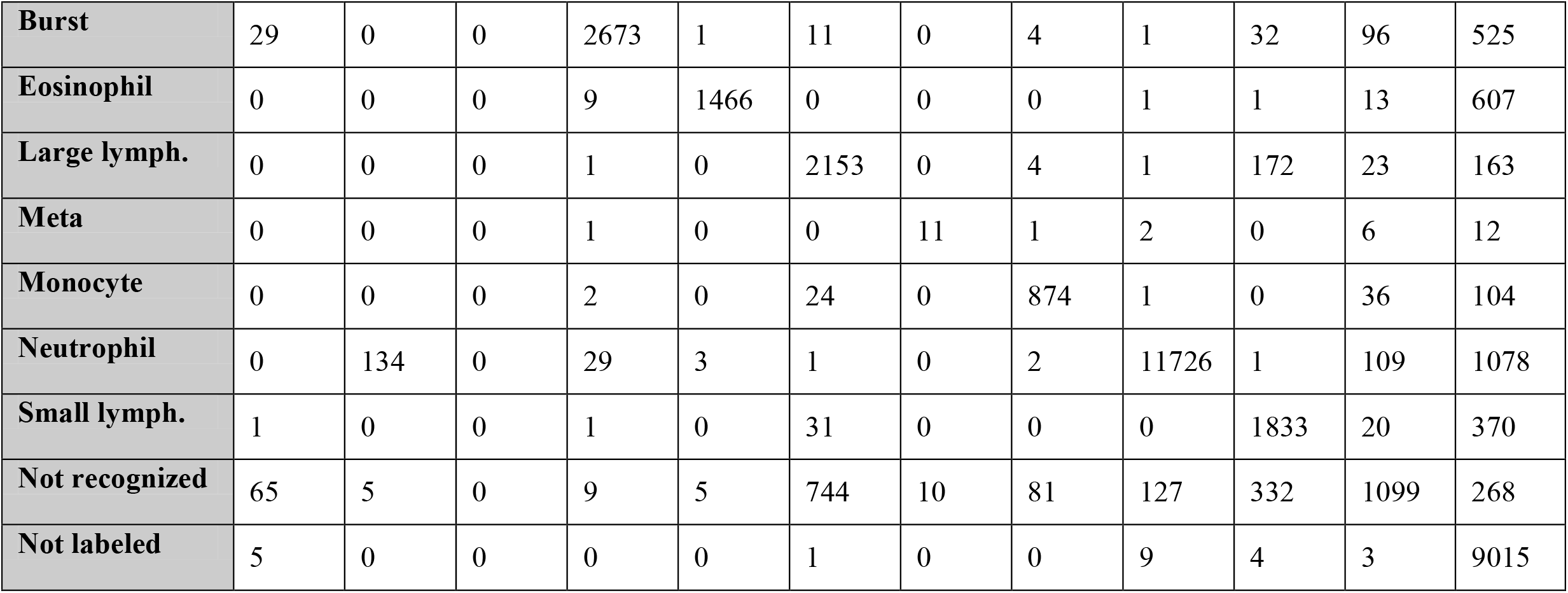
The number of labels associated with two experts. The rows and columns belong to the first and second experts, respectively.

#### Data structure

Raabin-WBC dataset consists of images that were taken from blood films (similar to figure 5). Corresponding to each microscopic image, a dictionary (.json format) file containing the following information about that image was provided:

- Information about the blood elements in the image including their coordinates and labels. Most of the elements are labeled by two experts.
- Information about the blood smears including staining method and the type of the disease. Note that all blood smears have been prepared from normal samples. Only a Chronic Myeloid Leukemia (CML) sample has been used to extract basophils.
- Information about the microscope includes the type of microscope and its magnification size.
- The type of the camera used.

There is also a subset of the database called double-labeled Raabin-WBC which includes cropped images of the five main types of WBCs and were labeled the same by both of the experts. We will explain more about this sub-dataset in the experiment section.

### Data Collection

The steps of data collection (figure 3) include preparing blood smears and photographing them, extracting the bounding box of white blood cells, data cleaning, and finally labeling the data and extracting ground truths. More details are explained in the rest of this section.

**Figure 3.**
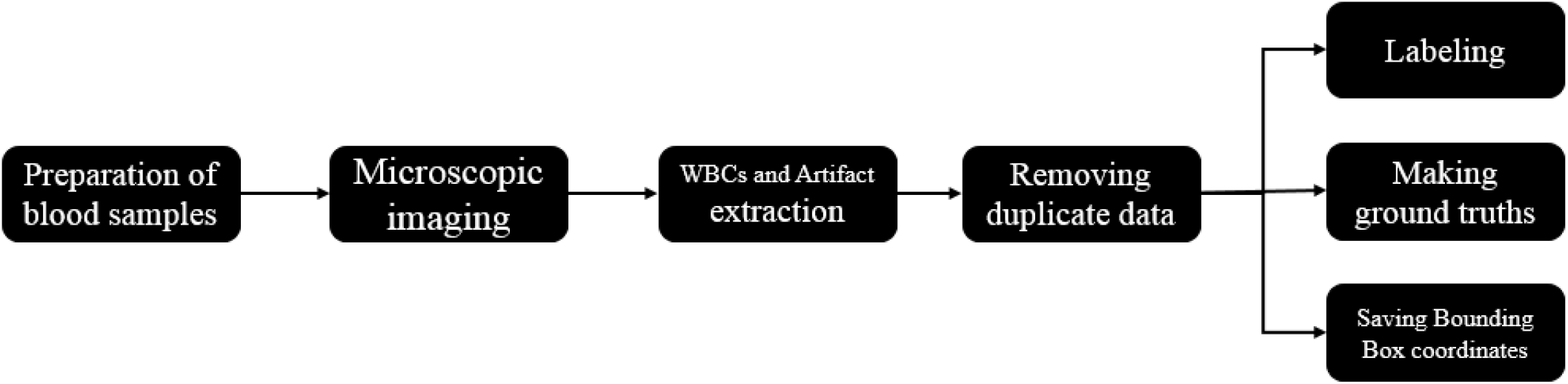
The main steps of Raabin-WBC dataset collection.

#### Preparation of blood smears and Imaging

72 normal peripheral blood films (male and female samples from ages 12 to 70) have been used to collect neutrophils, eosinophil, monocyte, and lymphocyte images. On the other hand, due to the very low presence of basophils in normal specimens (<1-2%) [6], basophils of one CML-positive sample have been imaged. Owing to the widespread use of Giemsa in medical labs [6], all samples were stained by Giemsa. Since there is no personal information about the sample donors, we were offered no chance to seek their permission for imaging. All samples were collected from the laboratory of Razi Hospital in Rasht, Gholhak Laboratory, Shahr-e-Qods Laboratory and Takht-e Tavous Laboratory in Tehran, Iran. Then, the process of imaging the slides was performed by the help of two microscopes called Olympus CX18 and Zeiss at a magnification of 100x. Since determining the Diff area to evaluate and count different types of white blood cells is of utmost importance, an expert lab staff had supervised the cell imaging process.

With the smart phones being widely used in the society, a rapidly growing trend has emerged with the aim of adapting them to medical diagnostics [21, 22]. The availability, ease of use and low cost of high-pixel density cameras available in smart phones make their imaging potential widely known in various science fields [23, 24, 25, 26, 27, 28, 29]. Therefore, in compiling this database, the cameras available on smart phones have been used, the details of which are given in Table 5. Smartphones can be adapted for microscopic imaging using some accessory equipment [30, 31, 32]. In order to prepare this database with the aim of facilitating the use of smart phones in microscopic imaging, an adapter was designed and made by 3D printing to mount the smart phone to the microscope ocular lens (figure 4). The designed adapter has somewhat managed to minimize the drawbacks of the commercial models available in the market such as restrictions on the size of the phone and ocular lenses, as well as the difficulty of the adjustment.

**Table 5.** Smartphone camera specifications used for data collecting.

| Smartphone | Release date | Sensor Model | Sensor Type | No. of Pixels | Aperture | Sensor Size | Pixel Size |
| --- | --- | --- | --- | --- | --- | --- | --- |
| **Samsung Galaxy S5** | 2014 | Samsung S5K2P2XX ISOCELL | CMOS | 16 MP | f/2.2<br>31mm | 1/2.6" | 1.12μm |
| **LG G3** | 2014 | Sony IMX135 Exmor RS | CMOS | 13 MP | f/2.4<br>29mm | 1/3" | 1.12μm |

**Figure 4.**
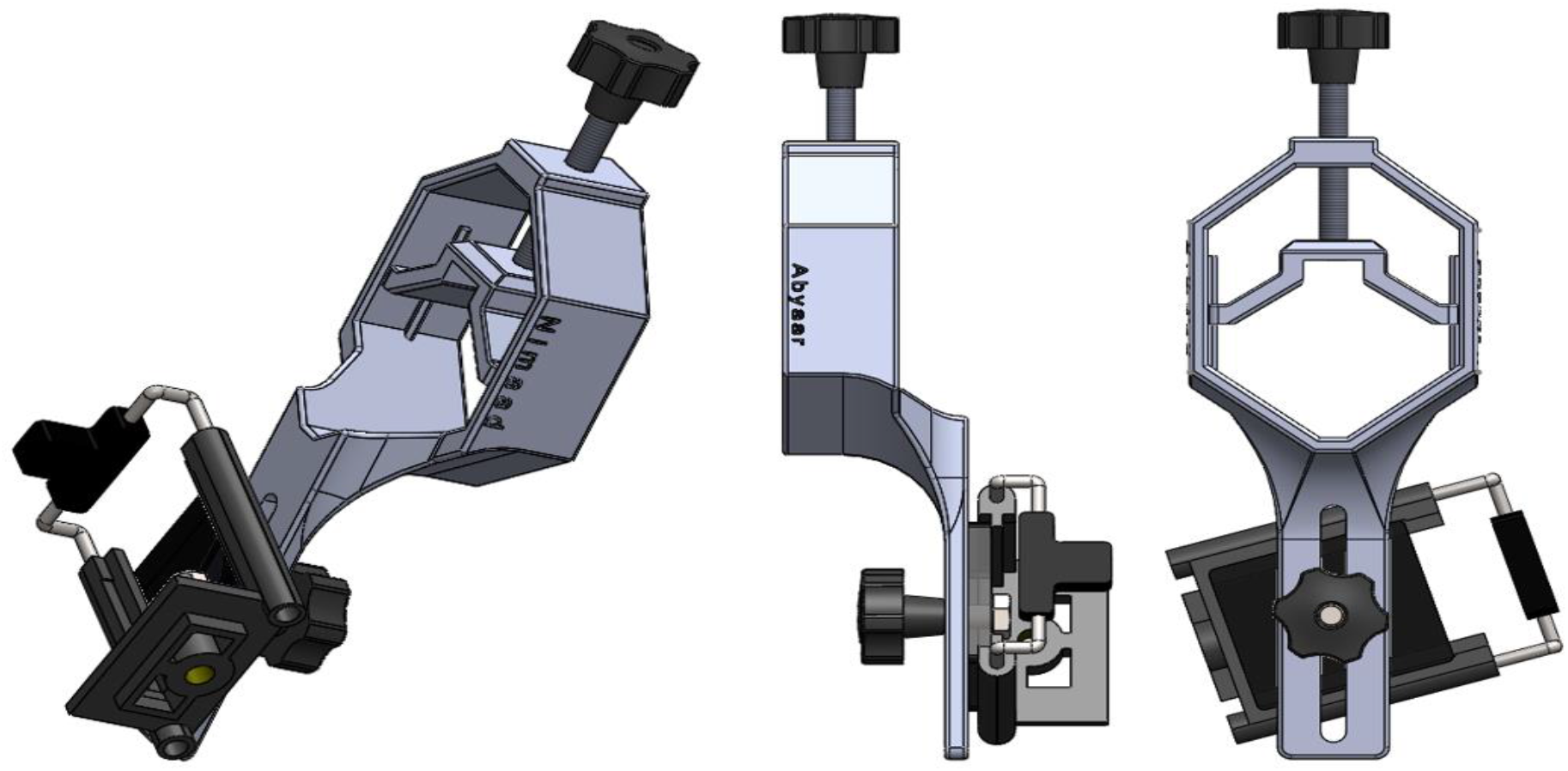
Designed adapter to mount smartphone to ocular lens of microscope.

#### Extraction of white blood cells from images

In total, about 23,000 images were taken from blood films. There exist many red blood cells in each blood smear image. It is also probable that one or more other blood elements such as white blood cells and sometimes color spots exist in the image. The bounding box of these blood elements should be somehow identified. For this purpose, two approaches have been considered. Due to the distinct color of the nucleus in white blood cells, in the first approach, a number of white blood cells were extracted manually as Grand Truth data, and a color filter was trained to separate the white blood cells from the background. The aforementioned color filter was applied to the main images, and the approximate position of the white blood cells was marked. Finally, a 512 by 512 square with the center of the cell is considered as a bounding box. In the second approach, extracted bounding boxes with the help of the first approach were used, and a Faster RCNN network [33], which is able to determine the exact location of the white blood cells in the original image, was trained. Eventually, about 43,000 blood elements were obtained.

#### Data cleaning

In the process of imaging from blood smears, a white blood cell may be placed in more than one image (figure 5); therefore, duplicate cell images exist among cropped images. The most important problem is that the two images of one cell are not necessarily very similar, and therefore, a simple mean square error on the value of the pixels is not enough to detect duplicate cell images. Indeed, a cell can be repeated more than twice. In figure 6, an example of three images of one cell is represented. As you can see the qualities of the three images are different.

**Figure 5.**
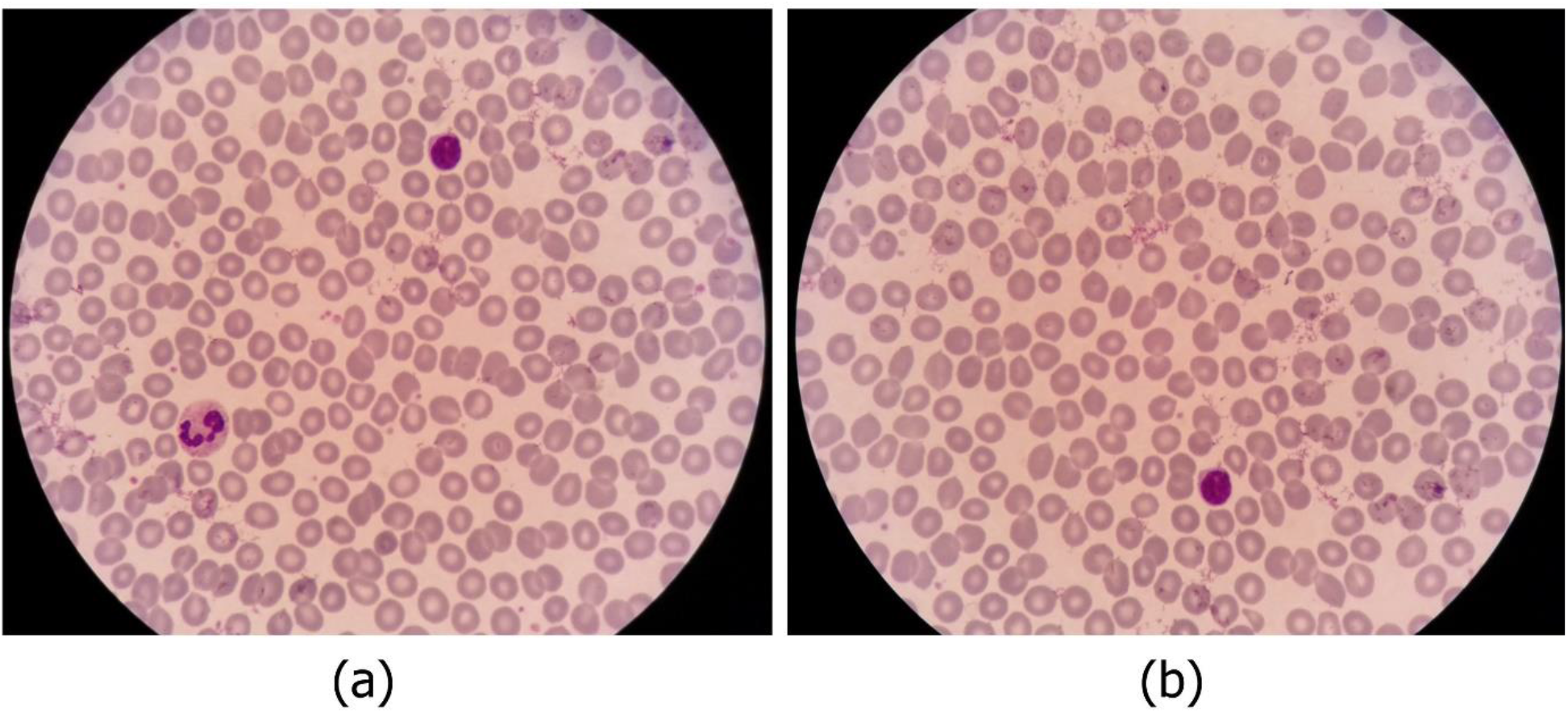
An example of two overlapped microscopic images.

**Figure 6.**
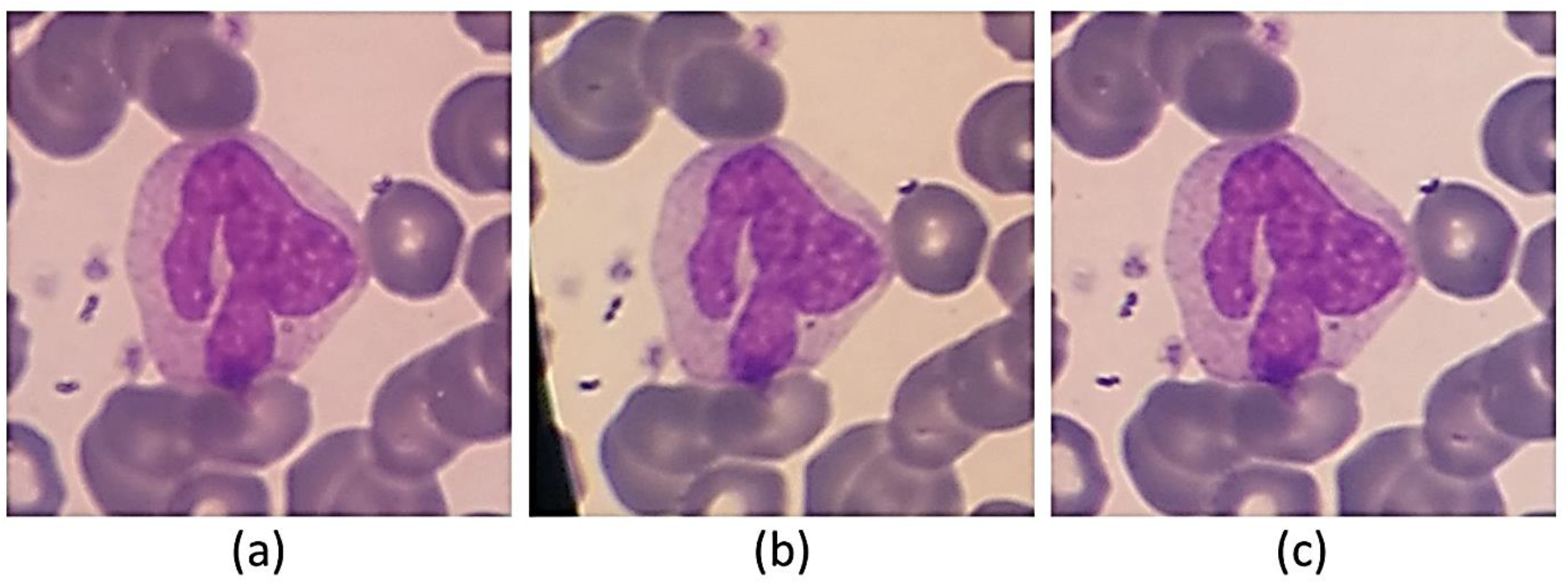
: One sample that had been repeated three times.

Manual comparison of these images in pairs is practically impossible. Hence, an artificial intelligence algorithm, fast and accurate, has been developed to remove duplicate cell images. We used Python ImageHash library, in this regard. First, for all pairs of cropped images, the Average Hash (AHash) and Perceptual Hash (PHash) values are calculated very quickly. Paired images, the AHash and PHash distances of which are less than those of the specific thresholds, are the same, and one of them should be removed. The thresholds of the Average Hash and Perceptual Hash are set manually through trial and error (See appendix1 for more details).

Since an image may exist more than twice, a two-by-two comparison is not sufficient. For this purpose, a solution to the problem is presented from Graph’s point of view. In fact, we have a graph with N nodes (N is the number of cropped cell images from blood film). There exist edges between the nodes that satisfy the sameness of condition. In this case, the connected components of the graph form equal images. Connected components of a graph can be calculated with the help of the breadth-first search algorithm very swiftly (See appendix2 for more details). If a connected component has n> 1 images, n-1 of them must be removed. To enhance the quality of the database, the image with the highest resolution remains out of n images, and the rest are deleted. The OpenCV [34] library is used to compare the image resolution. In this regard, Sobel horizontal and vertical filters [35] are applied to the images and the gradient magnitude is calculated for each pixel. Finally, the image with the highest average gradient magnitude is the sharpest one.

As described in section 2, to preserve more information, we prepared our data in the form of large images not in the form of cropped images (like figure 5). For each large image, the coordinate and the labels of the containing cells are provided. We tried to remove as many duplicates as possible from large images. Indeed, the large image in which all containing cells are inside the other images should be removed. For example, in figure 5, the image b should be removed.

### Labeling Process

This section describes the labeling process, which involves determining the cell types and the ground truth of the nucleus and cytoplasm. As you can see in table 1, the characteristics of nucleus and cytoplasm can significantly affect determining the type of the cell. Some papers [36, 16] extract different features from the nucleus and the cytoplasm to classify white blood cells. These features usually describe the shape and the color of the nucleus and the cytoplasm.

#### Cell type labeling

For labeling cells, two applications were developed for Android (Figure 7). One application is for labeling cropped cells (Figure 7-part b) and the other one is for selecting the location and type of the cells (Figure 7-part a). Furthermore, a desktop application with the help of Python Tkinter library [37] was developed in order to assign the location and type of the cells (Figure 8). It is worth mentioning that most of the images were labeled by two experts.

**Figure 7.**
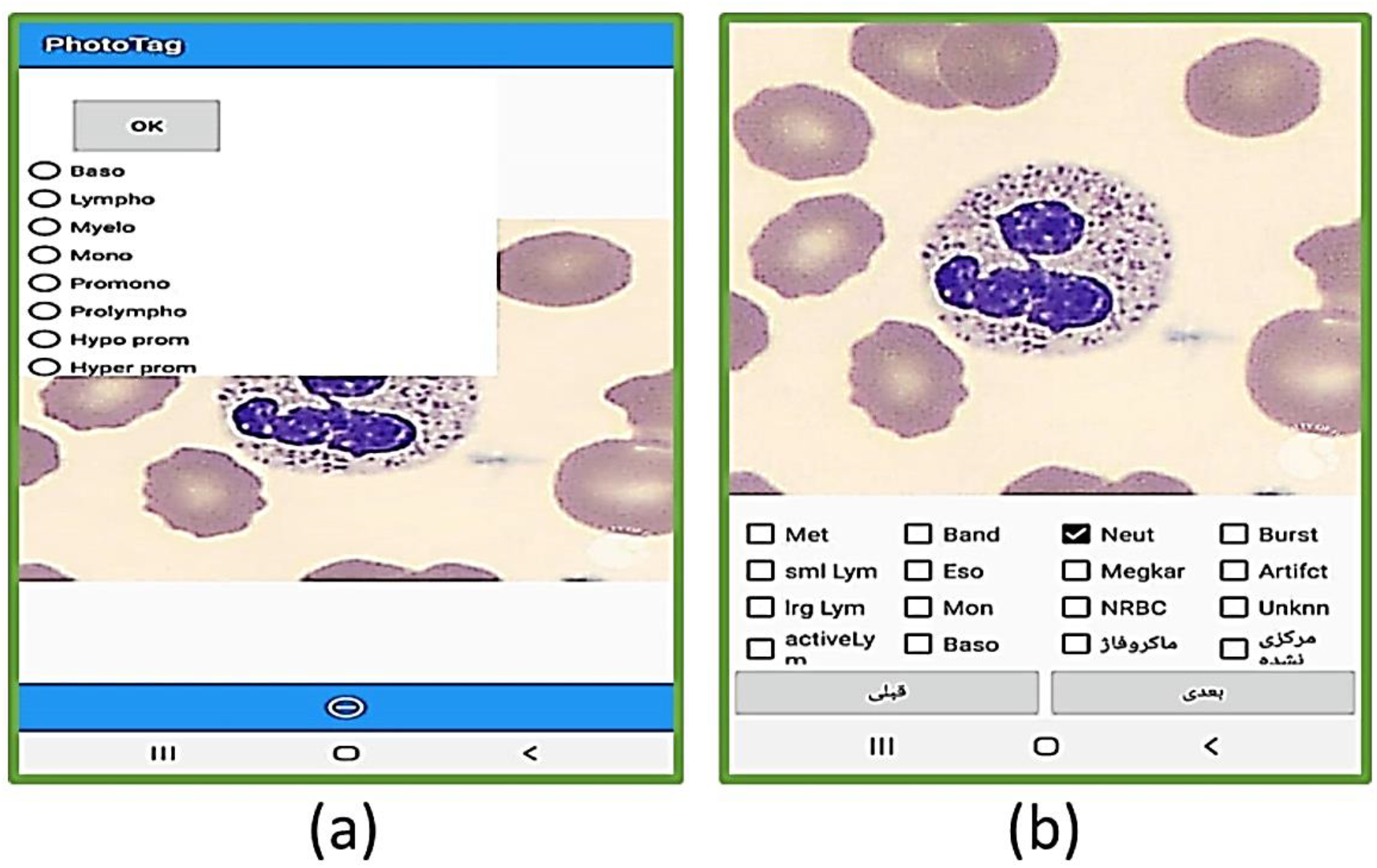
The user interface of the two android applications that were designed to selecting and labeling the white blood cells.

**Figure 8.**
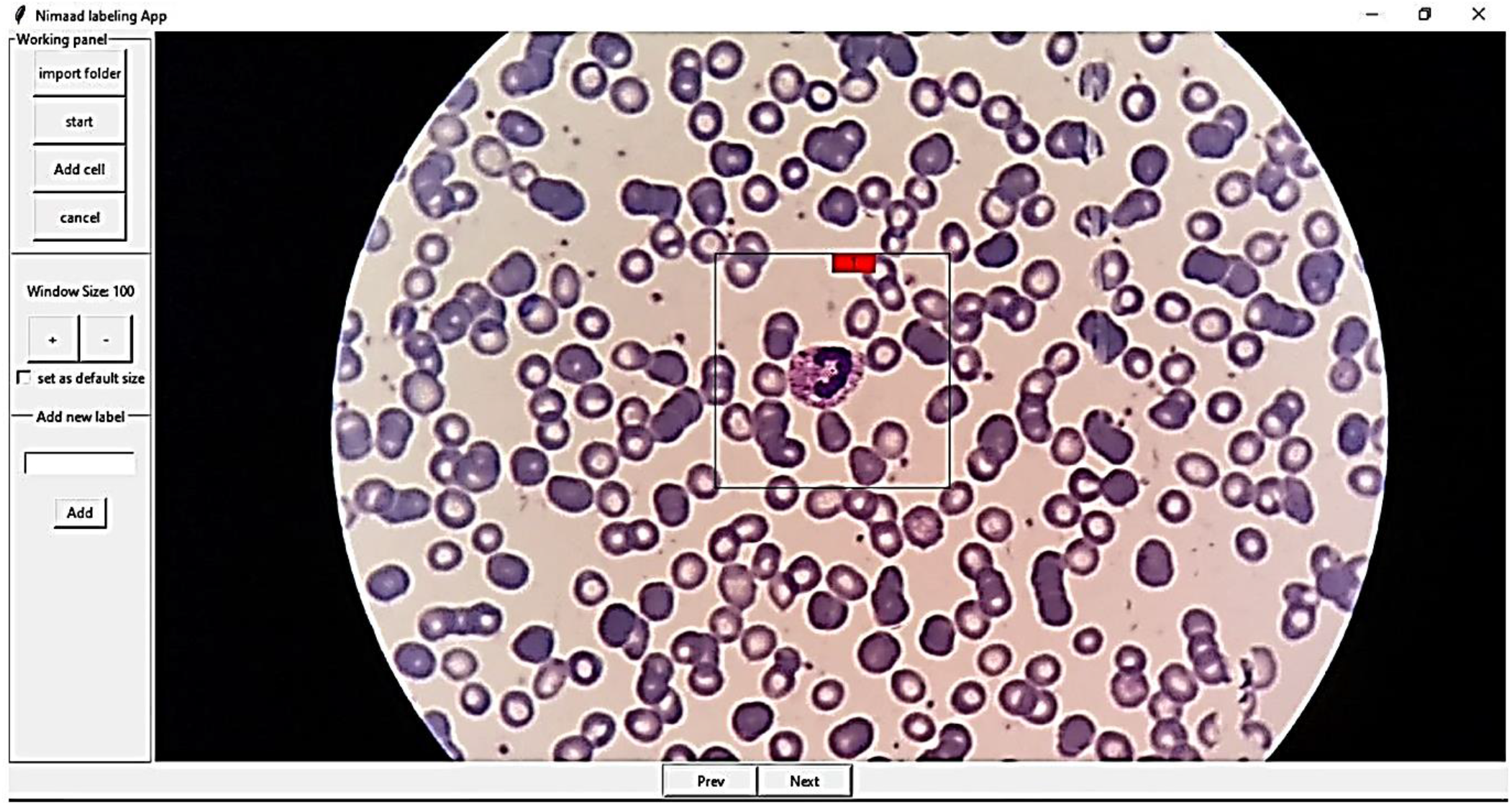
The user interface of the desktop application designed for labeling white blood cells.

#### Ground truth of the nucleus and the cytoplasm

In recent years, many researchers have investigated segmenting the cytoplasm and nucleus of the white blood cells [3, 15, 16, 17, 18, 20]. Hence, we tried to prepare the ground truths of the cytoplasm and the nucleus for a proper number of cropped white blood cells. For this purpose, 1145 cropped images including 242 lymphocytes, 242 monocytes, 242 neutrophils, 201 eosinophils, and 218 basophils were randomly selected, and their ground truths were extracted by an expert. It is worth mentioning that we only prepared the ground truth of the whole cell for basophils, and we were not able to produce the basophils’ cytoplasm and nucleus ground truth. This is because the basophils are usually covered by very purple granules, and the border between cytoplasm and nucleus is not easily obvious. Figure 10 shows some samples of the cells along with their ground truths.

To produce the nuclei’s ground truth, a new published software called Easy-GT [38] was employed. This software has been developed to extract the ground truth of the nucleus. In Easy-GT software, the nucleus is determined by a relatively accurate segmentation method, and if necessary, the user can adjust the ground truth of the nucleus by modifying the final threshold [38] (Figure 9). In the segmentation process, the RGB image is first color-balanced [38] and converted to CMYK color space. Secondly, the two-class Otsu’s thresholding algorithm [39] applied to the M channel gives us a threshold (*th*^2*class*^). Again, the three-class Otsu thresholding algorithm is applied to the M channel and the two lower and upper thresholds 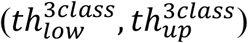 are extracted. Finally, the ultimate threshold value is obtained by computing the convex combination of *th*^2*class*^ and 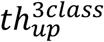.

**Figure 9.**
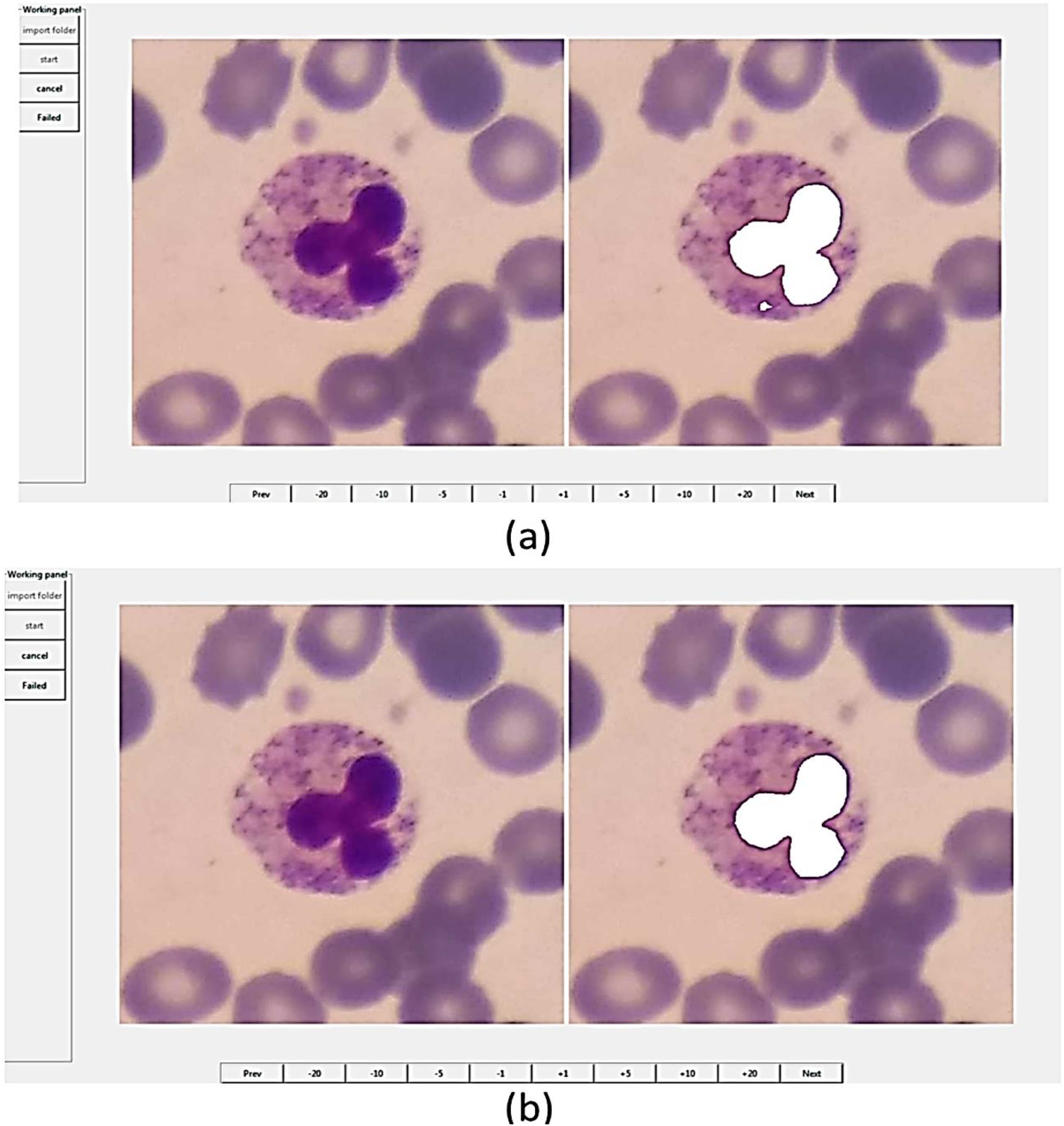
The user interface of Easy-GT software [38]. This software is developed in order to extraction of nucleus ground truth in white blood cells.

**Figure 10.**
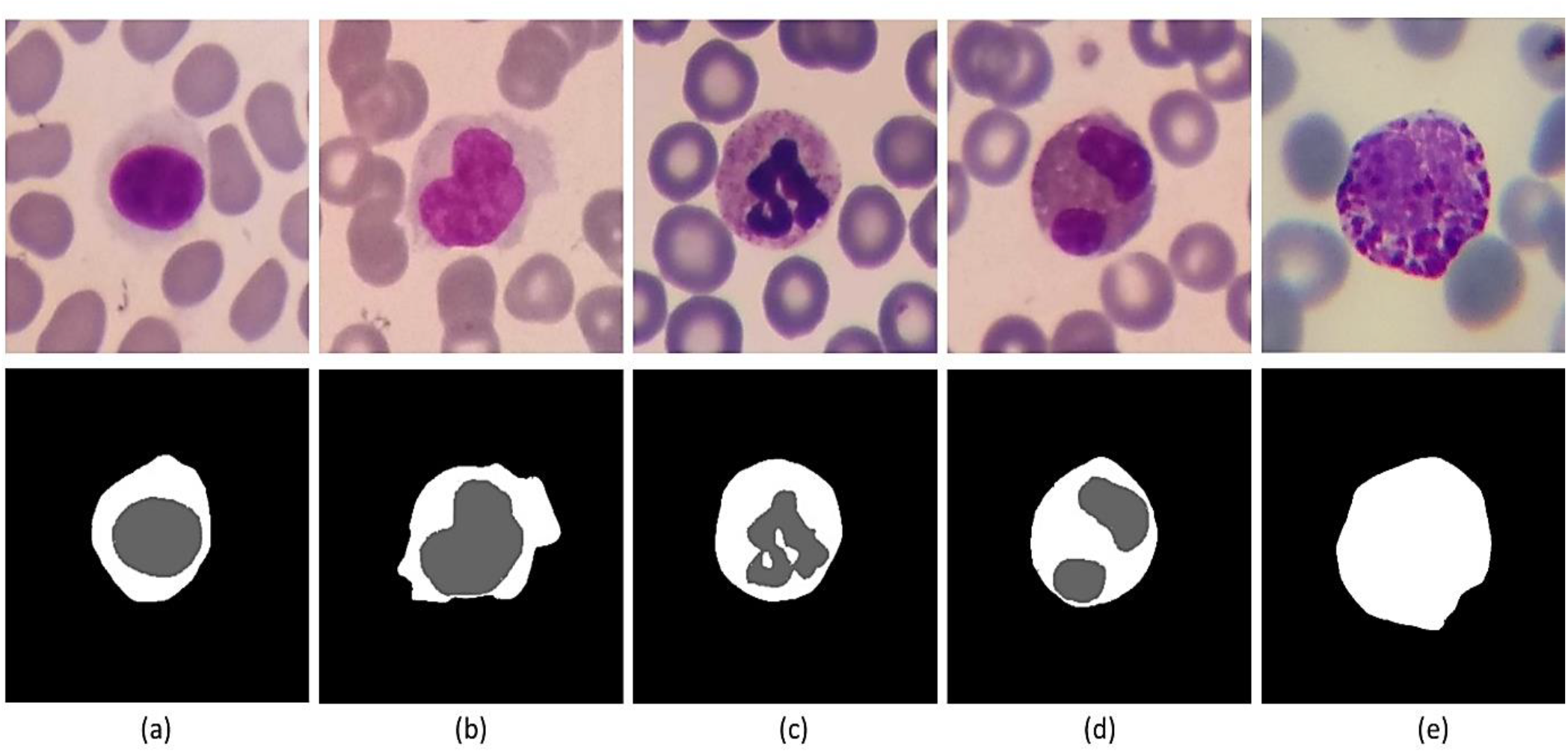
Some samples of ground truths that provided in Raabin-WBC dataset. First row is original cropped image of white blood cells. Second row is ground truths. (a), (b), (c), (d), and (e) are lymphocyte, monocyte, neutrophil, eosinophil, and basophil, respectively.

To make the ground truth of the cytoplasm, a light pen was used, and the ground truth of the whole cells were specified by an expert. Finally, by removing the nucleus part obtained from Easy-GT, only the cytoplasm remains.

### Experiments

In this section, we are going to do some machine learning experiments on Raabin-WBC data. Due to the diversity of information in the database, many research lines can be defined. Yet, we consider the most common possible experiment. We classify five classes of white blood cells, and we leave the rest to those who are interested in this field. For this purpose, we used the double-labeled cropped cells, and considered only five main classes including mature neutrophils, lymphocytes (small and large), eosinophils, monocytes, and basophils. We called this sub-dataset Double-labeled Raabin-WBC. In the following, we will compare this database with some existing 5-class databases and train some deep popular neural networks. We will also discuss the generalization power of the models.

#### A comparison with similar Datasets

Various datasets of normal peripheral blood with different properties exist, but in general, most of them have a small number of samples. This is due to the fact that in the medical field, data collection and labeling are complicated. On the other hand, in the field of Hematology, AI models are usually sensitive to some specifications of the dataset such as the number of data, the staining technique, the microscope and camera used, and the magnification. So, by altering the aforementioned characteristics, the accuracy of the models may be reduced. In table 6, the characteristics of some datasets have been presented and compared with Double-labeled Raabin-WBC. As you can see, our database is far better in several ways including data number, label assurance, Ground Truth, camera and microscope variety. Most importantly, this database is available to everyone for free.

**Table 6.**
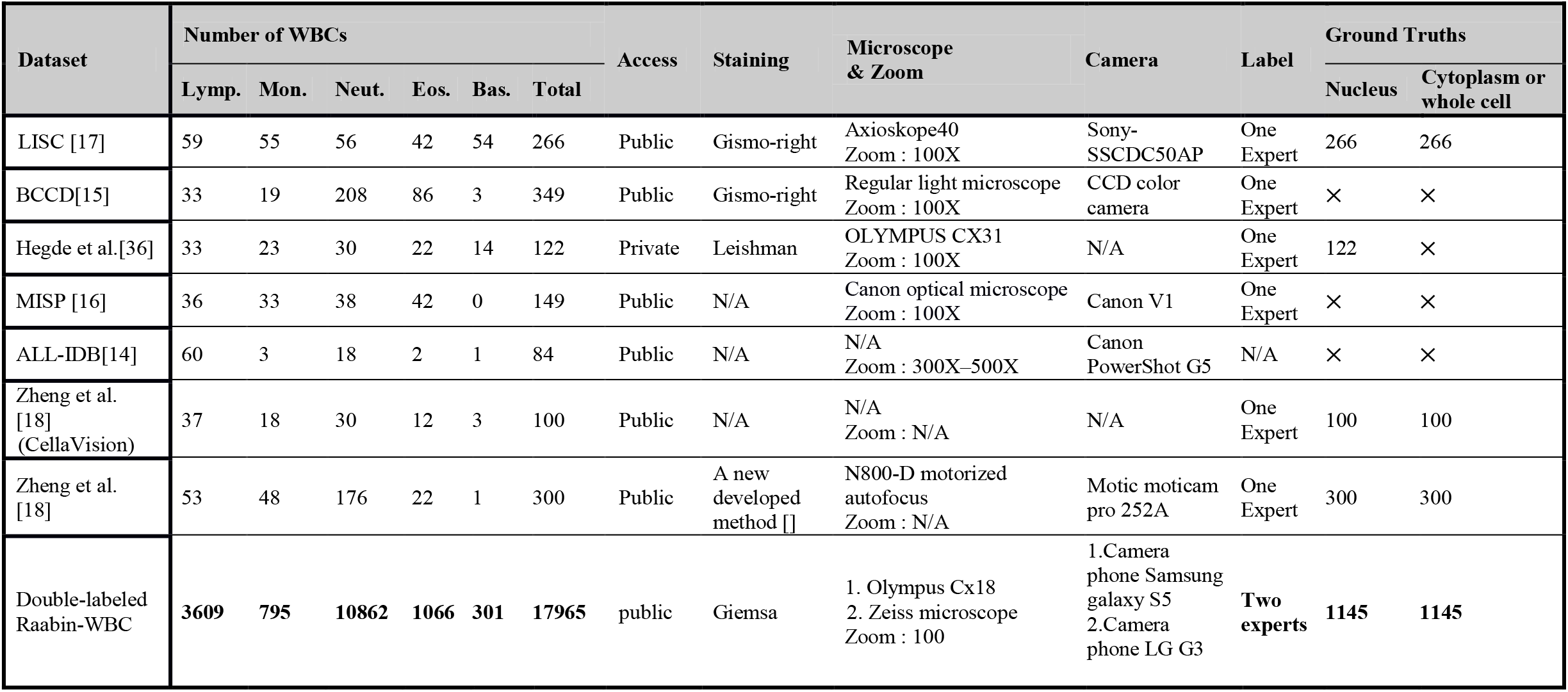
Comparing some datasets with double-labeled Raabin-WBC. Double-labeled Raabin-WBC does not contain the repeated samples as well as includes only five general cells (lymphocyte, monocyte, neutrophil, eosinophil, and basophil).

#### Utilized models

Some popular pre-trained deep neural networks have been trained on Double-labeled Raabin-WBC in order to classify five types of white blood cells. VGG16 [40] which is the oldest CNN used simply consists of alternating convolutional and pooling layers. From deep residual network families, Resnet18 [41], Resnet34 [41], Resnet50 [41], and Resnext50 [42] were tested. In Resnet architecture, the identity shortcut connections that skipped one or more layers are used [41]. Resnext is an extension of Resnet in which the residual block is replaced by a new aggregation component [42]. In mentioned aggregation component, the input feature map is projected to some lower-dimensional representations, and their outputs are combined by summing [42]. DenseNet121 [43] is another chosen CNN which consists of dense blocks. In each dense block, each layer is fed from all previous layers, and its outputs are transferred to all next layers.

Another used deep architecture is MobileNet-V2 [44] which is suitable for mobile devices. The building block of MobileNet-V2 is an inverted residual block, and non-linearities are removed from narrow layers. MnasNet1 [45] and ShuffleNet-V2 [46] are other light-weight CNNs for mobile devices. In MnasNet, reinforcement learning is employed to find an efficient architecture [45]. In ShuffleNet-V2 [46] at the beginning of the basic blocks, a split unit divides the input channels into two branches, and at the end of the block, concatenation and channel shuffling occur. Beside the aforementioned neural networks, we also utilized a feature-based method [47] in which the nucleus was segmented at first, and its convex hull was then obtained. After that, shape and color features were extracted using the segmented nucleus and its convex hull. Finally, WBCs were categorized by an SVM model.

#### Classification results

The generalization power of the models described in the former section is to be examined at two levels. For this purpose, we split data into three groups of training data, test-A, and test-B, the properties of which can be observed in table 7. The quality of the images in the test-A dataset is similar to that of the training dataset, but the images in the test-B dataset have different qualities in terms of camera type and microscope type. Unfortunately, the test-B data only contains double-labeled neutrophils and lymphocytes.

**Table 7.** The number of samples in Training data, Test-A, and Test-B.

| Sets | Lymph. | Mono. | Neut. | Eos. | Bas. |
| --- | --- | --- | --- | --- | --- |
| Training data | 2427 | 561 | 6231 | 744 | 212 |
| Test-A | 1034 | 234 | 2660 | 322 | 89 |
| Test-B | 148 | 0 | 1971 | 0 | 0 |

Since the training data are not balanced, in other words, the number of cells in each class is imbalanced. Hence, the training set was augmented and moderated using augmentation methods such as horizontal flip, vertical flip, rescaling, and a combination of them. In order to evaluate the models, four metrics are considered for each class: precision (P), sensitivity (S), F1-score, and accuracy (Acc). The aforementioned criteria are obtained through the equations (1), (2), (3), and (4).

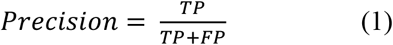

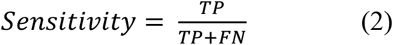

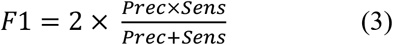

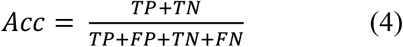

In equations 1–4, TP, FP, TN, and FN are true positive, false positive, true negative, and false negative, respectively. In tables 8 and 9, the results on test-A and test-B datasets are presented. Also, in the last row of the tables 8 and 9, the results of the feature-based classification presented in the paper [47] have been displayed. In figure 11 the plots of the accuracy and the loss of training data and validation data related to nine pre-trained models are shown.

**Table 8.** The results of the different pre-trained models as well as Tavakoli et al. [47]’s method on test-A dataset.

| Methods | Lymph |  |  | Mono |  |  | Neut |  |  | Eosi |  |  | Baso |  |  | Acc (%) |
| --- | --- | --- | --- | --- | --- | --- | --- | --- | --- | --- | --- | --- | --- | --- | --- | --- |
|  | P (%) | S (%) | F1 (%) | P (%) | S (%) | F1 (%) | P (%) | S (%) | F1 (%) | P (%) | S (%) | F1 (%) | P (%) | S (%) | F1 (%) |  |
| **ResNet18 [41]** | 98.84 | 99.23 | 99.08 | 96.96 | 95.30 | 96.12 | 99.66 | 99.36 | 99.5 | 95.77 | 98.45 | 97.09 | 100 | 100 | 100 | 99.06 |
| **ResNet34 [41]** | 98.66 | 99.52 | 99.09 | 96.93 | 94.44 | 95.67 | 99.85 | 99.14 | 99.49 | 94.67 | 99.38 | 96.97 | 100 | 100 | 100 | 99.01 |
| **ResNet50 [41]** | 98.47 | 99.52 | 98.99 | 97.36 | 94.44 | 95.88 | 99.74 | 99.40 | 99.57 | 96.94 | 98.45 | 97.69 | 100 | 100 | 100 | 99.10 |
| **ResNext50 [42]** | 98.01 | 100 | 98.99 | 99.53 | 91.45 | 95.32 | 99.74 | 99.55 | 99.64 | 97.85 | 98.76 | 98.30 | 100 | 100 | 100 | 99.17 |
| **MnasNet1 [45]** | 98.93 | 98.65 | 98.79 | 94.14 | 96.15 | 95.14 | 99.66 | 98.76 | 99.21 | 92.15 | 98.45 | 95.20 | 100 | 100 | 100 | 98.59 |
| **MobileNet-V2 [44]** | 98.85 | 99.32 | 99.08 | 96.93 | 94.44 | 95.67 | 99.66 | 99.40 | 99.53 | 96.97 | 99.38 | 98.16 | 100 | 100 | 100 | 99.12 |
| **DenseNet121 [43]** | 98.38 | 99.61 | 98.99 | 97.72 | 91.45 | 94.48 | 99.70 | 99.14 | 99.42 | 94.40 | 99.38 | 96.82 | 100 | 100 | 100 | 98.87 |
| **ShuffleNet-V2 [46]** | 98.09 | 99.32 | 98.70 | 96.49 | 94.02 | 95.24 | 99.74 | 99.36 | 99.55 | 97.85 | 98.76 | 98.30 | 100 | 100 | 100 | 99.03 |
| **VGG16 [40]** | **98.26** | **98.26** | **98.26** | **95.09** | **91.03** | **93.01** | **99.43** | **98.57** | **99** | **89.01** | **98.14** | **93.35** | **100** | **100** | **100** | **98.09** |
| **Tavakoli et al. [47]** | **97.23** | **95.07** | **96.14** | **84.87** | **86.32** | **85.59** | **98** | **95.60** | **96.78** | **72.24** | **91.30** | **80.66** | **96.59** | **95.51** | **96.05** | **94.65** |

**Table 9.** The results of the different pre-trained models as well as Tavakoli et al. [47]’s method on test-B dataset.

| Method | Lymph. |  |  | Neut. |  |  | Acc (%) |
| --- | --- | --- | --- | --- | --- | --- | --- |
|  | P (%) | S (%) | F1 (%) | P (%) | S (%) | F1 (%) |  |
| ResNet18 [41] | 21 | 94 | 34 | 100 | 2 | 3 | 8 |
| ResNet34 [41] | 24 | 94 | 38 | 100 | 27 | 43 | 32 |
| ResNet50 [41] | 21 | 95 | 35 | 100 | 2 | 4 | 8 |
| ResNext50 [42] | 24 | 90 | 38 | 100 | 3 | 5 | 9 |
| MnasNet1 [45] | 20 | 88 | 33 | 100 | 0 | 0 | 6 |
| MobileNet-V2 [44] | 72 | 50 | 59 | 100 | 0 | 0 | 4 |
| DenseNet121 [43] | 43 | 64 | 51 | 100 | 8 | 15 | 12 |
| ShuffleNet-V2 [46] | 42 | 75 | 54 | 100 | 0 | 0 | 5 |
| VGG16 [40] | **96** | **89** | **93** | **100** | **65** | **79** | **66** |
| Tavakoli et al. [47] | **94** | **54** | **69** | **100** | **92** | **96** | **90** |

**Figure 9.**
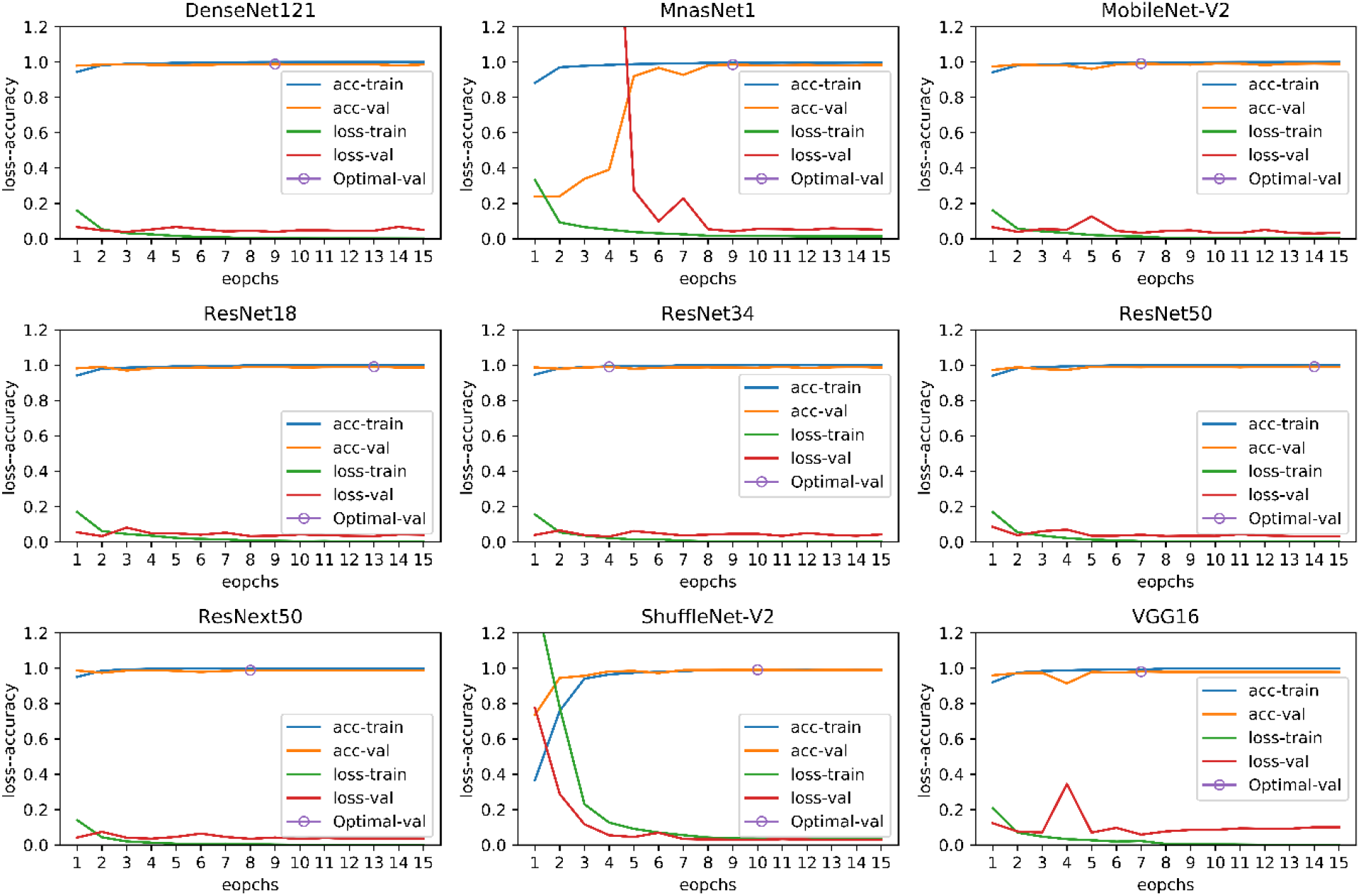
The plots of the accuracy and loss of training data and validation data related to nine pre-trained models.

The results are surprising, and all methods have an acceptable outcome on test-A data. Yet, the performance of most of the models on test-B data has experienced a dramatic decrease. The feature-based method [47] had the least performance reduction, despite having the lowest accuracy on the test-A data. Among deep neural networks, the VGG16 [40] network has relatively more generalizability. It can be said that the feature-based method could extract more meaningful features from cell images than the deep neural networks. If we had not tested the models on test-B data, we would have thought that we have trained a strong classification model; yet, this was not the case. In this experiment, we do not want to conclude that deep neural networks have less generalization power than feature-based methods. If we applied some appropriate pre-processing on the images before training or used some smarter image augmentation methods, the performance of deep neural networks would be better. In this experiment, you can easily understand the role of the dataset in the training of machine learning models.

All training processes were carried out using a single NVIDIA GeForce RTX 2080 Ti graphic card and were handled by Python 3.6.9 and Pytorch library version 1.5.1. We considered 15 epochs for the training process and the starting learning rate, and the batch size were 0.001 and 10, respectively. The learning rate was decayed by the ratio of 0.1 and step size 7. Stochastic gradient decent was utilized as an optimization method. We used torchvision library in order to load pre-trained networks on the ImageNet dataset [48]. The output size of the last linear layer was changed from 1000 to 5.

## Conclusion

By evaluating the peripheral white blood cells, a wide range of benign diseases such as anemia and malignant ones such as leukemia can be detected. On the other hand, early detection of some of these abnormalities, such as acute lymphoid leukemia, despite its lethality, can help its treatment process. Therefore, it is important to adopt methods that can be effective in early detection of the disease. The role of machine learning methods in intelligent medical diagnostics is becoming more and more prominent these days. Indeed, deep neural networks are revolutionizing the medical diagnosis process and are considered as one of the stare-of-the-arts.

Since deep neural networks usually have a huge number of training parameters, the overfitting problem is not highly unlikely. Therefore, the diversity of training data is necessary and cannot be ignored. In medical diagnostics, in particular, this diversity gets bolder, for the medical devices can be very diverse. For example, in the field of hematology, the type of microscope and camera is very influential. To this end, we tried to collect a huge free available dataset of white blood cells from normal peripheral blood so as to relatively satisfy the mentioned diversity. This multipurpose dataset can serve as a reference dataset for the evaluation of different machine learning tasks such as classification, detection, segmentation, and localization.

## Acknowledgments

The authors would like to thank the laboratory of Razi Hospital in Rasht, Gholhak laboratory, Shahr-e-Qods laboratory and Takht Tavous laboratory in Tehran for their efforts in collecting samples of the dataset. Ms. Anahita Ahouraei and Mr.Arman Yekta are also appreciated for the efforts they put in editing Raabin database contract as our attorneys at law. The authors would like to acknowledge the financial support of University of Tehran Science and Technology Park for this research under grant number 130067, as well.

## The Availability of Data

The Raabin-WBC dataset is publicly available through the following link: https://www.raabindata.com/free-data/

## Appendix 1

In this section, we aim at explaining the image hash algorithms that we used for removing the duplicate images. The Average hash (A-hash) and Perceptual hash (P-hash) were employed to evaluate the similarity of the two cells. The Average hash has a very simple algorithm including the following steps:

- The image is resized to 8*8 pixels and converted to grayscale.
- The mean value of the pixels is computed: The pixels above this value take the value of 1, and the others take the value of zero. Indeed, we have a binary sequence with a length of 64 in row major order, for instance.
- The final hash is computed by converting the binary sequence to base 64.

The average hash is very fast and also robust against changes in brightness, contrast, color, size, and aspects of ratio. The perceptual hash which is more robust than the average hash includes the following steps:

- The image is resized to 32*32 pixels and converted to grayscale.
- The discrete cosine transform is computed and resized to 8*8 pixels.
- The mean value of the pixels is computed and similar to the average hash a binary sequence is obtained.
- The binary sequence is converted to base 64.

In order to compare two images, first, the desired hash is computed for each image. Then, the hamming distance between two hashes should be computed. After doing some experiments, we concluded that using the average and perceptual hash simultaneously provides better performance. We set two thresholds for the average and perceptual hash through trial and error (11 for A-hash and 14 for P-hash). Paired images with A-hash and P-hash distances less than tuned thresholds, are the same, and one of them should be removed.

## Appendix 2

In this section we will explain more about the graph theory topics we used for removing duplicate images. Consider we have an undirected graph *G*(*V, E*) that *V* and *E* are the sets of vertices and edges, respectively. A connected component is a subgraph *G*′ (*V*′ ⊆ *V, E*′ ⊆ *E*) in which there is a path between all pairs of nodes in *V*′. Both depth-first search (DFS) and breadth-first search (BFS) can be used to find all connected components of the graph [1]. We used BFS for this purpose. The pseudocode of the algorithm are as follows:

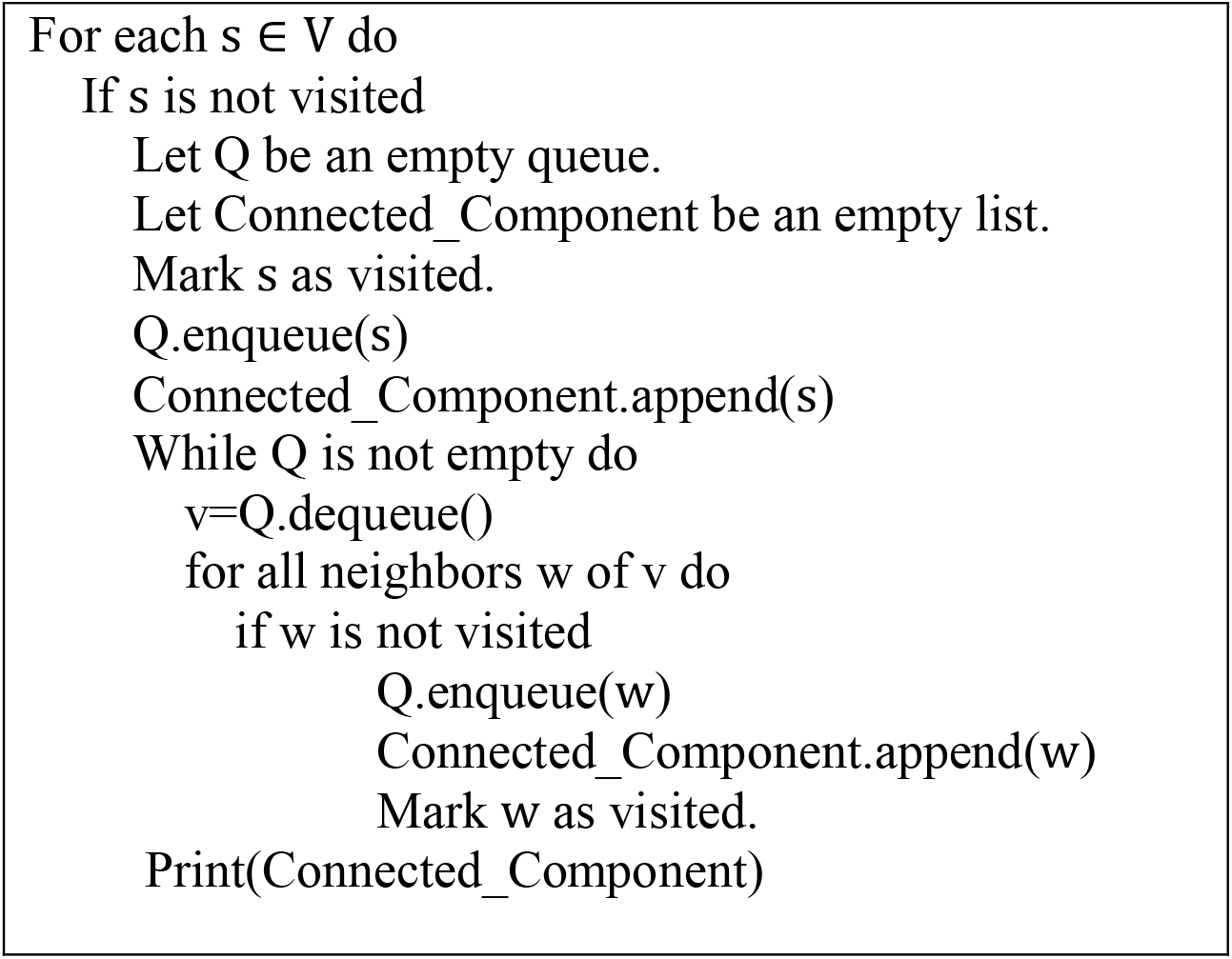

## Author contributions

Z. M. K. is the main author and the supervisor and executor of the artificial intelligence and image processing parts. S. S. is also the main author and the head of the medical and data collection department. E. T. and P. R. helped to manuscript revision and data collection. M. A., F. M., E. S. S., M. G., F. G., and S. M. have somehow helped in the data collection and labeling process. R. H. is the advisor of the team and the corresponding author.

## Additional Information

**Correspondence** and requests for materials should be addressed to R. H.

